# Porous marine snow differentially benefits chemotactic, motile, and non-motile bacteria

**DOI:** 10.1101/2022.05.13.491378

**Authors:** Benedict Borer, Irene Zhang, Amy E. Baker, George A. O’Toole, Andrew R. Babbin

## Abstract

Particulate organic carbon settling through the marine water column is a key process that regulates global climate by sequestering atmospheric carbon. The initial colonization of marine particles by heterotrophic bacteria represents the first step in recycling this carbon back to inorganic constituents – setting the magnitude of vertical carbon transport to the abyss. Here, we demonstrate experimentally that bacterial motility is required for particle colonization and chemotaxis specifically benefits at higher settling velocities. We further explore the role of particle microstructure on the colonization efficiency of bacteria with different motility traits. We highlight that non-motile cells benefit disproportionally from the porous microstructure and are relatively enriched in the particle wake due to the efficient particle colonization of chemotactic and motile cells. Our results imply that although the chemotactic and motile bacteria benefit from the high nutrient availability when colonizing the particles, scavenging of these cells benefits the often oligotrophic, non-motile cells common among the planktonic community.

**Significance statement:** Bacteria in the ocean rely on ephemeral nutrient patches from sinking marine particles, but attaching to these structures is challenging as particle settling rates often exceed bacterial swimming velocities and the numerically dominant marine bacteria are non-motile – posing an interesting paradox about the prominence of particle foraging. Here, we quantify the importance of chemotaxis and motility for the efficient colonization of marine particles and find that although chemotaxis provides a clear advantage, motility is the basic requirement for particle colonization. We expand this analysis to consider highly heterogeneous particle structures and find a disproportionate benefit for non-motile cells by facilitating a direct encounter with the particle surface and enriching non-motile microbes in the nutrient-rich particle plume.

## Introduction/Narrative

Heterotrophic bacteria exert a fundamental role on ocean biogeochemistry by recycling dissolved and particulate organic matter back to inorganic constituents. The most abundant bacterial species in the marine environment are non-motile^1^ and rely on diffusive fluxes to consume the fairly homogeneous but low concentrations of dissolved and recalcitrant carbon in the bulk water^2^. In contrast, particulate organic matter such as phytoplankton aggregates or fecal pellets (together termed “marine snow”) sink through the water column, resulting in a vertical flux of carbon from the surface mixed layer to the abyss. This process of sequestering atmospheric carbon through photosynthesis and subsequent transport to the ocean interior and sediments is termed the biological carbon pump, and is a key driver in regulating Earth’s global carbon cycle^3^. Compared to the surrounding bulk water, these particles can have orders of magnitude higher carbon and nutrient concentrations and act as microbial hotspots in an otherwise nutrient poor marine environment^4^. The intensified metabolic activity within sinking particles^4,5^ can drive rapid remineralization of the organic matter during settling and ultimately determine the efficiency of the organic carbon pump^6,7^. It has been shown that the particle associated microbial community is distinct when compared to the surrounding, free living community^8,9^ and is typically enriched in fast-growing copiotrophic bacteria^10–12^ associated with motile and chemotactic behavior.

Marine particles exist across a spectrum of shapes and sizes^13^. The fate of these particles is governed by their remineralization dynamics and settling velocity, which themselves are complex functions determined by their size, shape and excess density (difference in average particle density compared to surrounding seawater)^14^. Across the globe, particle size distributions are captured by a power law, suggesting that large particles (e.g., >3 mm) are rare whereas the numerically dominant particles are very small (<1 mm). As a result, some large particles (namely, fecal pellets) may settle at extremely fast rates up to a few hundred meters per day^14^ that play an important role in carbon sequestration as it limits the time for remineralization by heterotrophic bacteria. Nevertheless, the average particle in the ocean typically sinks very slowly^15^, ranging between 1 and 10 m d^-1^. Yet, at the scale of a bacterial cell, particles settling at seemingly slow velocities may still outpace the swimming velocity of a bacterium. For instance, a modest settling velocity of 5 m d^-1^ translates to 58 µm s^-1^ and — especially when considering the stochastic random walk patterns frequently observed in flagellated bacterial species^16^ — presents a scenario where many bacterial species cannot effectively chase fleeting particles. Therefore, colonization of marine particles must occur in a very short window of time during the bacteria-particle encounter when the particle passes by the bacterial cell. Indeed, bacterial cells need some form of self-propulsion to traverse the steep velocity gradients in close proximity of the particle surface during colonization^17^. Furthermore, the physical shape of the bacterial cells (e.g., from cocci to rods and filaments) can govern the colonization efficiency through shear-driven reorientation of their trajectory^17,18^.

Many motile bacterial cells have optimized their swimming behavior by directing their movement in response to a chemical stimulus, a phenomenon known as chemotaxis. Peering into mechanisms of how chemotaxis governs bacterial life in the ocean is challenging^19^ yet experimental systems that quantify a chemotactic response are insightful. For instance, milli- and microfluidic devices have been used to quantify prevalence and importance of chemotaxis *in situ*^*20*,*21*^ and to study the optimal foraging of bacteria on dynamically emerging synthetic nutrient patches *in vitro*^*22*^. Furthermore, localized nutrient patches resulting from phytoplankton exudation, copepod death and fecal pellets have been used to investigate the chemotactic capabilities of marine cells towards biogenic attractants^23,24^. However, the noted experimental systems study chemotaxis in an absence of fluid flow which results in a homogeneous diffusion field around the nutrient hotspots and therefore do not include the additional constraint of the advective flow field. For settling particles, the flow around the particle warps the diffusive field into a nutrient rich plume^25^ where chemotactic bacteria can navigate the resulting spatiotemporal gradients^26^ and exploit these short-lived but important nutrient rich patches^27^. An experimental system to investigate if and how chemotaxis facilitates colonization of particle surfaces in the presence of a continuous carbon source leaking from the particle under the influence of flow is necessary.

In this study, we use a millifluidic experimental system and fluorescence microscopy (Fig. 1) to quantify how bacterial motility and/or chemotaxis facilitate the colonization of synthetic marine particles that maintain a constant nutrient source (representative of leaky marine snow). By employing fluorescently tagged mutants that are either chemotactic, non-chemotactic but motile, or nonmotile, we can differentiate the contribution of different motility traits to particle colonization across controlled flow conditions mimicking different settling velocities. We combine these experimental observations with computational predictions based on solving coupled Navier-Stokes and reaction-diffusion equations with bacterial cells represented as individual agents to explore how colonization efficiency varies with bacterial traits. We investigate how the micro-structure of more realistic, porous marine snow (resulting from aggregation of biogenic polymers and inorganic minerals or an organic particle being actively degraded) affects bacterial colonization, and furthermore, compare the porous particles with their rough-surface and smooth-surface counterparts. In addition, there exists a seeming paradox: if motility and chemotaxis are so beneficial in supporting heterotrophs to find food, why are non-motile cells by far the most numerically dominant in the ocean? We highlight how marine particles may act as microscale sponges that scavenge the chemotactic and motile cells from the surface ocean and create a top-down control that facilitates the non-motile cells to dominate the planktonic microbial community.

**Figure 1:**
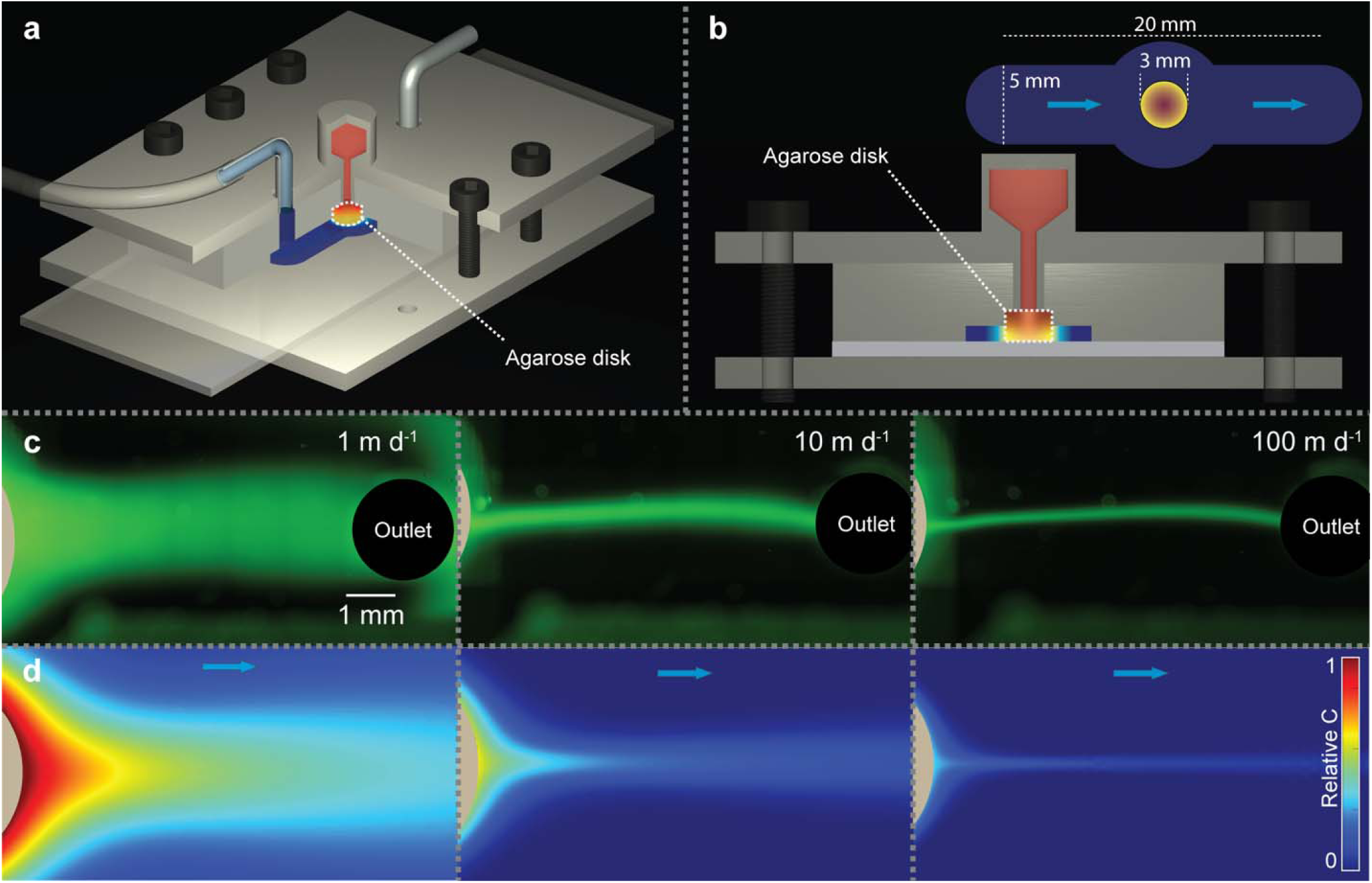
Conceptual image of the experimental system and resulting nutrient plume. a) The experimental system includes a millifluidic device (Polydimethylsiloxane - PDMS) on top of a glass microscope slide with a synthetic marine particle in the form of a 3 mm diameter agarose disk. Nutrients leak from the sterile reservoir (red) through the agarose disk into the flow chamber (dark blue) and mimics a constantly leaking particle. The flow chamber is sealed to the glass slide using two 3D printed parts that are screwed together with the nutrient reservoir directly engineered inside the upper part. The synthetic marine particle that connects the two systems ensures a steady leak of nutrients from the agarose disk into the main flow chamber. b) The total length of the flow chamber is 20 mm with a channel width of 5 mm. The diameter of the central expansion is 8 mm to accommodate the 3 mm agarose particle and keep the channel width constant at 5 mm across the whole channel length. The flow channel is 1 mm in depth with the agarose disk embedded and additional 1 mm within the PDMS for structural stability (i.e. a total height of 2 mm). c) We imitate different settling velocities from 1 to 100 m d^-1^ using a syringe pump and visualize the resulting particle plume using fluorescein. d) We use a digital twin approach (congruent geometry, flow and nutrient conditions as in the experiments) to simulate the resulting flow pattern and particle plumes that are in good agreement with the experimental observations (panel c).

## Methods

### Bacterial strains and culture conditions

We performed the experiments with a motile and chemotactic wild type *Pseudomonas aeruginosa* PA14, a chemotaxis deficient mutant *P. aeruginosa* PA14 *cheZ:*:*TnM* (deficient in dephosphorylation of the CheY protein which transmits sensory signals to the flagella motors^28^) and a motility deficient mutant *P. aeruginosa P*A14 Δ*fliC* (deficient in production of flagellin, the primary component of the flagellum^29^). We confirmed the fluorescence and phenotype of the mutants using fluorescence microscopy and motility/chemotaxis assays (SI Fig. 1). All strains were routinely kept in 25% glycerol at -80°C and grown in Lennox Lysogeny broth (5 g L^-1^ NaCl, BD Life Sciences, further abbreviated LB) medium or on 1.5% agarose LB plates. All strains were tagged with the same yellow fluorescent protein (pUC18T-mini-Tn7T-Tp-eyfp, Addgene plasmid #65035) following the mini-TN7 insertion protocol^30^. After conjugation, 300 µg ml^-1^ Trimethoprim (Trp) was used for routine growth and maintenance of all strains to ensure pure cultures.

### Measurement of fluorescence and bacterial density in liquid culture

We used a Tecan Spark plate-reading fluorometer and spectrophotometer (Tecan Trading AG, Switzerland) to verify the monotonic and increasing relationship between the optical density of a liquid bacterial culture at 600 nm (further referred to as OD600) and their fluorescent intensity for all three strains (SI Fig. 2a). We grew overnight cultures of the three strains in LB after which we diluted each culture to an OD_600_ of 1. We then created a serial dilution (four technical replicates, five dilution steps) by diluting 20% of the culture in 80% fresh LB medium. We transferred 250 µL of the final diluted cultures into a 96 well plate and added two rows of controls containing sterile LB medium. We subsequently measured the absorbance at 600 nm and fluorescent intensity at 513 nm excitation and 530 nm emission wavelengths characteristic of eYFP for all wells.

**Figure 2:**
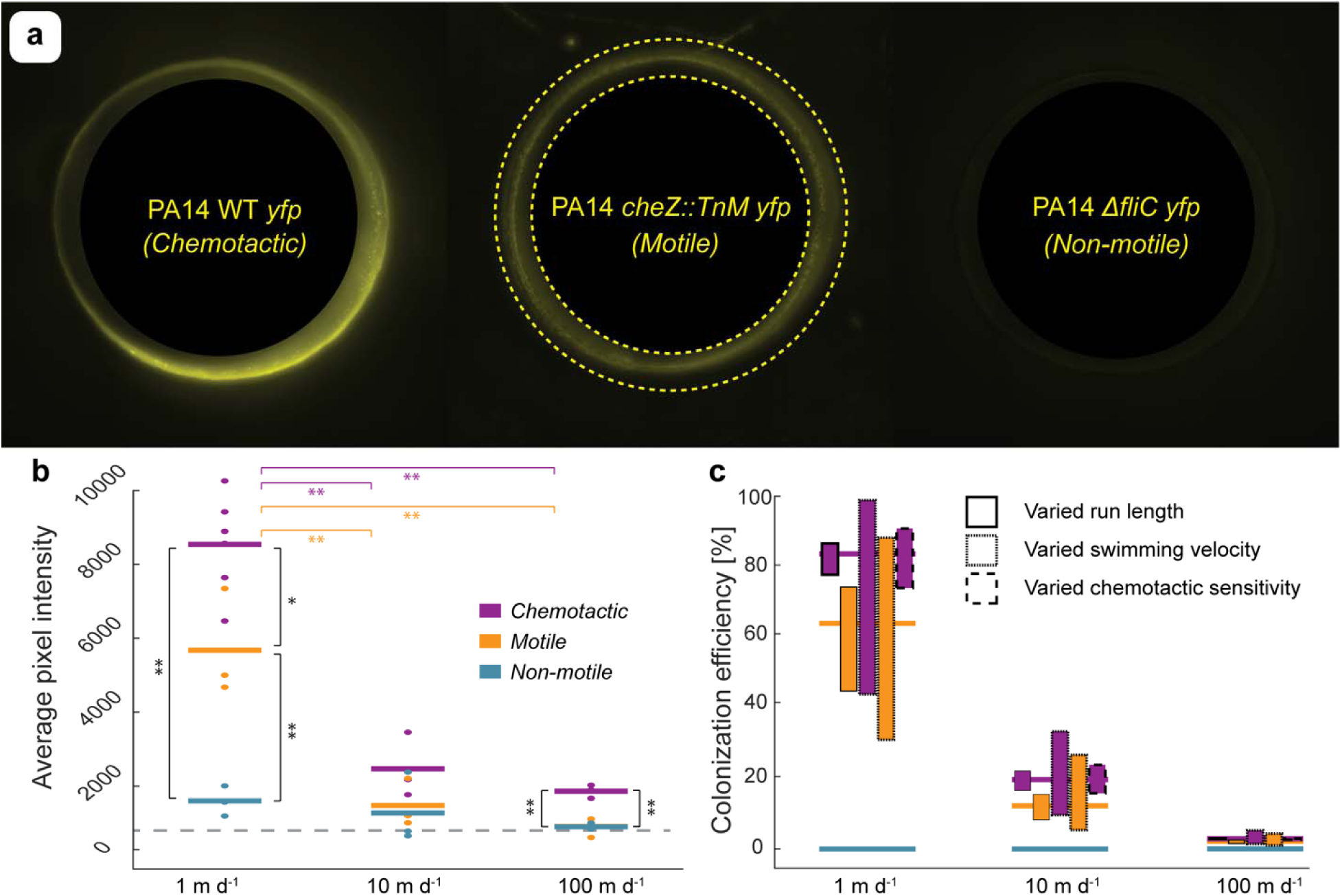
Experimental colonization results and comparison to mathematical predictions. a) We use the fluorescence intensity of the three mutant strains as a proxy for the number of cells that colonized the particle surface within a region of ±250 µm from the particle surface (delineated by the yellow dashed circles in the central image). These results show colonization for a simulated settling velocity of 10 m day^-1^ with a flow direction from left to right. b) Quantification of the fluorescence signal as a proxy for the total number of colonized cells with pairwise comparisons between the strains for the same settling velocities and within the same strain across different settling velocities (* for p<0.05, ** for p<0.01). The dashed gray line indicates the mean fluorescent intensity of sterile particles. c) Prediction of colonization using the numerical model where the horizontal lines show the mean value across 10 replicates using the standard parameterization of bacterial motility and bars show the variation when strategically exploring the sensitivity across the parameter space in the chemotaxis and motility algorithm (Methods, Fig. 3).

### Fabrication of millifluidic devices

We fabricated the millifluidic devices using Poly(dimethylsiloxane) (PDMS) and 3D printed molds. All 3D devices were printed using a Form3 stereolithography printer (Formlabs, Somerville, MA, USA) and clear resin (RS-F2-GPCL-04) at 25 µm resolution with Form Wash (20 minutes) and Form Cure (1 h at 60 °C) for post processing. We exposed the mixed PDMS (Polydimethylsiloxane, DOW, CAS# 68083-19-2, 10:1 ratio of base to curing agent) to vacuum for 30 minutes at room temperature before pouring it into the 3D printed molds. We subsequently exposed the molds a second time to vacuum for 30 min at room temperature before curing at 70 °C for 24 h.

### Colonization experiment using the leaky particle device

We grew the bacterial strains overnight in culture tubes containing 5 mL of LB medium (37 °C, 220 rpm). We then diluted 200 µL of the overnight cultures with 20 mL fresh LB medium and regrew the cultures (37 °C, 220 rpm) for 3h into exponential phase (typically reaching an OD600 between 0.5 and 1). We further diluted the cells to an OD_600_ of 0.1 and washed the cells twice by carefully centrifuging the cells at 1000 × *g* for 10 minutes and subsequently resuspending the cells in a saline medium void of nutrients^27^ (further called motility medium, containing 10 mM potassium phosphate monobasic (Sigma-Aldrich, CAS# 7778-77-0), 100 µM EDTA (Sigma-Aldrich, CAS# 6381-92-6), 10 mM NaCl (Sigma-Aldrich, CAS# 7647-14-5) and adjusted to pH 7.5 using 1M NaOH) by gently shaking for 5 minutes on an orbital shaker at 100 rpm. We then added 1 mL of this culture to 10 mL of motility medium, resulting in approximately 500,000 cells per mL, and transferred the final bacterial solution into a sterile syringe (Beckton-Dickinson, 10 mL slip tip). We verified that this procedure does not affect swimming capacity or shear off flagella.

We prepared synthetic marine particles using low melting agarose as previously described by Smriga et al. 2021^31^. In short, we dissolved 15 mg of low melting agarose (Agarose low gelling temperature, Sigma-Aldrich, CAS# 39346-81-1) in 1 mL of motility medium that contained 10 µM each of L-serine and L-aspartate in a block heater at 80 °C. After cooling for 3 minutes, we cast a 2 mm thick slab of agarose using a 3 mL syringe (BD, 3 mL Luer lock) and a 22G blunt needle (Industrial Dispensing Supplies) in between two microscope slides (VWR, Vistavision Microscope Slide, 75 mm x 50 mm x 1 mm) separated by 2 mm using additional glass slides. After solidifying at room temperature for 15 minutes, we punched the agarose discs from the agarose slab using 3 mm diameter biopsy punches (Integra Miltex, Disposable Biopsy Punch, 3 mm) which we transferred into the experimental device previously filled with motility medium containing the two chemoattractant amino acids in the reservoirs (L-serine and L-aspartate). We verified that the bacterial cells are unable to grow in the motility medium when supplemented with the two amino acids (SI Fig. 2b), confirming that observed increase in fluorescent intensity is due to bacterial attachment alone and not growth during the experiment.

### Fluorescent microscopy and image analysis

We used a Nikon Ti2-E inverted microscope equipped with an Andor Zyla 4.2 sCMOS camera and a SOLA SEii 365 Light Engine to visualize the colonized particles. The microscope was controlled using Nikon Elements (v.4). All images were captured in 16-bit with a 10x Plan Fluor DLL objective and an eYFP filter set for visualizing the bacterial cells (Chroma 49003, ex 500/20 nm, em 535/30 nm). Each particle was visualized at 5 Z levels with 50 µm between the layers and the center located approximately at the middle of the channel (i.e., ∼500 µm from the bottom glass slide which we located using the perfect focus system of the Nikon microscope). Each Z layer was captured as a 4 × 4 tile scan with a 20% overlap that was automatically stitched by the Nikon Elements software using the built-in blending algorithm. Individual images of the tile scans were captured at 50% illumination with a shutter speed of 800 ms for all particles and strains to ensure equal illumination of each particle and thus reliably quantify the colonized population. These settings resulted in an optimal dynamic range within the images for all strains and conditions tested and thus require the least amount of post processing.

All image analysis was performed in MATLAB R2021a (The Mathworks, Natick, MA, USA) using ND2Reader^32^ to import the ND2 files produced by Nikon Elements. For each particle, we only considered an area ±250 µm around the particle edge for quantification. Fluorescence intensities in Fig. 2 are reported as mean pixel values across the whole area surrounding the particles without any further post processing to minimize the potential bias from image postprocessing.

### Individual-based model of bacterial cells colonizing marine particles

We developed a mathematical framework to validate experimental results and gain further insights by combining computational fluid dynamics with reaction-diffusion modeling and an agent-based representation of motile bacterial cells. We used the FEATool Multiphysics software 1.15.2 (Precise Simulation Ltd.) in Matlab to predict the flow and nutrient field around the marine particles and a custom algorithm to simulate non-motile, motile and chemotactic cells. Here, we first describe the individual based representation of bacterial cells (which is identical for all geometries) and subsequently the specific geometries used for the different studies. The Navier-Stokes equations in combination with reaction-diffusion modeling was solved on a triangular mesh with a non-uniform resolution of 100 µm increasing to 10 µm at any boundary of the simulation for the experimental twin scenario and 1 mm increasing to 10 µm for the large rectangular domain. Example geometries and grid meshes are shown in SI Fig. 3.

**Figure 3:**
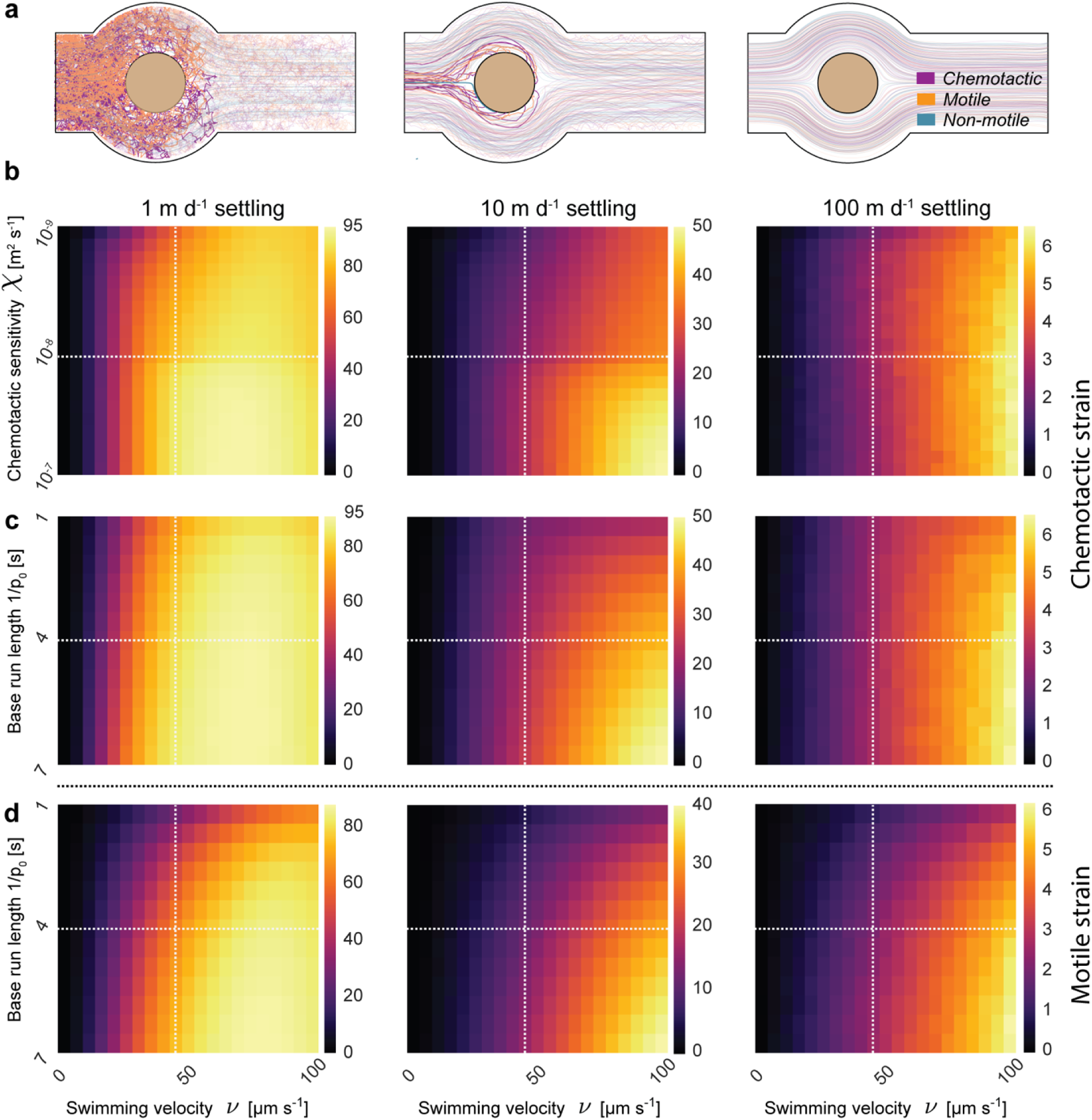
Sensitivity analysis of the motility and chemotaxis algorithms. a) Simulated trajectory of individual bacterial cells at different settling velocities. Highlighted trajectories represent cells that colonized the particle in contrast to faded trajectories that are not successful. The ability of bacterial cells to influence their trajectory with self-propulsion becomes negligible at rapid particle settling velocities. b) Colonization efficiency at different combinations of chemotactic sensitivity and cell swimming velocity for the chemotactic strain. c) Colonization efficiency at different combinations of base run length and cell swimming velocity for the chemotactic strain. d) Colonization efficiency at different combinations of base run length and cell swimming velocity for the motile strain. Dotted lines in panels b, c and d represent the parameter values for *P. aeruginosa* in the experimental twin geometries and stochastic simulations. We used different scales (shown by the color bar) for the different settling velocities with markedly lower colonization at higher settling velocities for all parameter combinations.

### Numerical simulation of bacterial motility

For all conditions and each replicate, we inoculate bacterial cells 6.5 mm upstream of the particle equator (i.e., 5 mm upstream of the leading edge of a 3 mm particle) uniformly random over a width of 4 mm. Each replicate simulates 100,000 cells of a single strain which results in an approximate inoculation density of 25 cells µm^-1^ across the channel. The model includes three modes of bacterial cell displacement: advective displacement by the fluid motion, active self-propulsion using a run and tumble mechanism, and random displacement due to Brownian motion. The displacement vector for each individual bacterial cell is determined by the local fluid flow field multiplied by the time step. All bacterial cells are additionally displaced by Brownian motion where the displacement vector is determined by the characteristic distance of diffusion for a particle of 1 µm diameter with the diffusion coefficient calculated using the Stokes-Einstein equation (*D* = *kT/6πηr*_*bac*_, with k the Boltzmann constant, T the temperature, *η* the dynamic viscosity and r_bac_ the bacterial cell radius). At 20 °C and a bacterial radius of 0.5 µm, each cell is displaced in a random direction by 92 nm in each numerical time-step of 0.01 s. When simulating motile cells, bacterial cells additionally use self-propulsion at a constant velocity in a fixed direction. Bacterial cells choose a new direction at random with a probability of *p*_0_ = 1/*τ* where p_0_ is the average tumbling probability per unit time [-] and *τ* the average run length of the bacterial cells [s]. The total displacement of bacterial cells in one time step is then the sum of all displacement vectors. Chemotactic cells have the ability to reduce their tumbling frequency as a function of positive nutrient gradients^33^, a feature which we capture in the model using the following Equation 1:

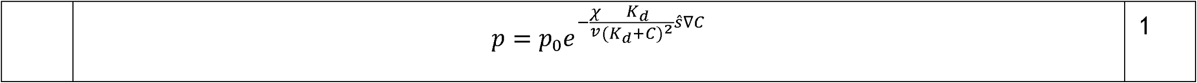

Where p is the chemotactically adjusted tumbling probability [-], p_0_ the tumbling probability associated with the average run length *τ* [-], *χ* the chemotactic sensitivity [m^2^ s^-1^], *v* the bacterial swimming velocity [m s^-1^], K_d_ the receptor/ligand dissociation equilibrium constant [mol m^-3^], C the nutrient concentration [mol m^-3^], ŝ the normalized vector of swimming direction [m] and v the nutrient gradient [mol m^-4^]. For positive nutrient gradients, Equation 1 describes an exponential decay of the tumbling probability. When experiencing a negative nutrient gradient (i.e., the cell is swimming away from the particle or the plume), we set the tumbling probability to p_0_ ^33,34^. The nutrient gradient is calculated based on the experienced change in nutrient concentrations over the past 0.3 s^35^ where the current nutrient concentration is taken from the nutrient concentration at the bacterial location. If the cell tumbles, a new direction is chosen at random. Finally, if cells collide with the *in silico* marine particle, we assume a “perfect sticking” condition (i.e., if the bacterial trajectory intersects with the particle boundary within a time step, we assume an irreversible attachment). We selected a time step of 10 ms which equates to a maximum advective travelling distance of 14 µm per time step at the highest settling velocity (which is below the smallest grid meshing and thus restricts numerical inaccuracies from the advection of cells). We set the total simulation time such that individual bacteria are advected a total of 60 mm even for the lowest flow of 1 m d^-1^ settling velocity.

### Experimental twin and no-boundary geometries

We mimic the experimental system using a digital twin approach. The domain has a total length and width of 20 mm and 5 mm, respectively, where a 3 mm particle is embedded in the center of an 8 mm spherical expansion in the center of the system to ensure the same channel width for the flow around the particle (to minimize flow acceleration and deceleration around the particle). Flow velocity is set as a parabolic profile at the inlet 10 mm upstream of the particle equator with a maximum flow velocity equal to the experimental condition (1, 10 and 100 m d^-1^). Due to the narrow channel (5 mm), bacterial cells are reflected by the boundary of the domain in case of collision whilst assuming a “perfect sticking” condition upon collision with the particle. We set a constant relative concentration boundary at the particle surface (C = 1) with a diffusion coefficient of 10^−10^ m^2^ s^-1^ that is representative of amino acids^36^. We furthermore create a large rectangular domain (50 mm × 50 mm) with a 3 mm particle at the center to investigate the influence of the constrained geometry in the experimental twin condition on the local flow field around the particle, associated nutrient plume and colonization probability, and better link our experiments to the real ocean.

### Porous particle geometry

Finally, we numerically simulate a cohort of 500 porous particles including their rough and smooth alterations to strategically investigate the influence of porosity, roughness and size on the colonization efficiency. Each particle is assigned a particle size at random between 100 µm and 3 mm and a target porosity between 0 and 0.7 (uniform distribution). We calculate a target number of sub-particles that form the aggregate using an average sub-particle diameter of 60 µm. We then create a stochastic geometry for each particle by choosing a location within the particle at random with a diameter between 20 µm and 100 µm for each of the sub-particles. The rough alteration of each particle includes a solid circular core with a diameter equal to the outer most sub-particle center and the smooth particle solely being this solid core (without any roughness). We calculate the resulting particle porosity using the particle surface area and total particle area. Each particle is located in a 5 cm by 5 cm domain where the settling velocity is assigned by Equation 2:

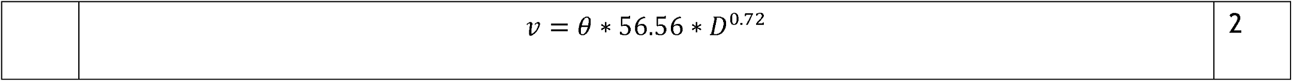

With v the settling velocity [m d^-1^], *θ* the particle porosity [-] and ESD the particle diameter [mm]. We set a constant relative nutrient concentration of 1 for each sub-particle boundary.

## Results

### Colonization of synthetic marine particles across motility traits

We use three genetically engineered strains of *Pseudomonas aeruginosa* PA14 in our experiments that constitutively express the same yellow fluorescent protein. A wild-type strain that is both motile and chemotactic (“chemotactic”), a chemotactic deficient mutant *P. aeruginosa* PA14 *cheZ:*:*TnM* that can nonetheless swim (“motile”) and a motility deficient mutant *P. aeruginosa* PA14 Δ*fliC* (“non-motile”). Motility and chemotaxis traits were confirmed by plate assay (SI Fig. 1). By using mutants of the otherwise isogenic parent strain, we can isolate the influence of chemotaxis and/or motility on colonization and directly compare the fluorescence signals captured with the microscope (Fig. 2a,b). Due to the inherent intricacies of this system and potential to bias the results through parameter choice, we refrained from converting the fluorescence signal to total cell number and only present the quantified fluorescent signal which is directly proportional to cell abundance (SI Fig. 2a). We observe and quantify the colonization of the particles by bacteria with different motility traits across three settling velocities that represent nearly suspended particles (1 m d^-1^), reasonable settling velocities for the majority of sinking particles (10 m d^-1^) and rapidly sinking, large particles that are numerically rare in the ocean but contribute overwhelmingly to the mass flux of the biological carbon pump (100 m d^-1^)^37^. For all settling velocities, we observe a consistent pattern where the chemotactic strain shows the highest mean fluorescence intensity, followed by the motile strain and lastly the non-motile strain (Fig. 2b, statistical significance in SI Table 1). In addition, the observed fluorescent intensity decreases for both the chemotactic and motile strains with increasing settling velocity (Fig. 2b). However, this decrease is less pronounced for the chemotactic strain compared to the motile strain (53%, 80% and 138% higher fluorescent intensity than the motile complements for 1 m d^-1^, 10 m d^- 1^ and 100 m d^-1^), suggesting that the chemotactic strain gains a competitive advantage over non-chemotactic motile cells at higher settling velocities due to the ability to rapidly respond during the ephemeral particle encounter. For the non-motile strain, we observed no significant difference in fluorescent signal across settling velocities.

Simulations show a markedly similar pattern of colonization efficiency when comparing the strains within and across the different settling velocities (Fig 2c). Here and throughout the manuscript, the colonization efficiency is defined as the fraction of the inoculated cells that ultimately attached to the particle. The high number of simulated cells (average density of 25 cells µm^-1^ across the channel at the inoculation location) results in negligible variance among the 10 technical replicates. For this reason, we only visualized the mean value of the simulation replicates in Fig. 2c (horizontal bars) and superimposed the variation in colonization efficiency when exploring the parameter space of the motility and chemotaxis algorithms (further described below). All pairwise statistical tests between the different strains at the same settling velocity for the simulation replicates were significant with the exception between the chemotactic and motile at a settling velocity of 100 m d^-1^. Congruent to the experimental results, both the chemotactic and motile strains showed a decrease in colonization efficiency with increasing settling velocities (p<0.01). The model predicts the highest benefit of chemotaxis at intermediate settling velocities with an increase in colonized cells of 31%, 60% and 34% for 1 m d^-1^, 10 m d^-1^ and 100 m d^-1^, respectively.

### Rapid swimming is key for particle colonization

We further explore the importance of the different motility and chemotaxis parameters for colonizing the particles (Fig. 3). Overall, the simulated settling velocity and associated localized flow velocity governs the ability of bacteria to influence their trajectory and colonization capability (Fig. 3a). The intensity of the response increases with settling velocity as the greater flow around the particle warps the nutrient concentration field into steeper nutrient gradients. Particle velocity profoundly influences the importance of chemotactic sensitivity (Fig. 3b). Interestingly, for both the slow and rapid settling velocities (1 m d^-1^ and 100 m d^-1^), the resulting colonization efficiency is primarily a function of the swimming velocity while the chemotactic sensitivity only contributes marginally. In contrast, the chemotactic sensitivity governs the colonization efficiency at an intermediate settling efficiency, particularly for cells with higher swimming velocities. A similar pattern to the one described above emerges when comparing the interaction between the swimming velocity and the base run length of the chemotactic strain (Fig. 3c). These data suggest that the reduction in base run length is compensated by the reduction in tumbling probability via chemotaxis when colonizing the particles. This concept is further supported when comparing the colonization pattern of the chemotactic strain to that of the motile strain (Fig. 3d). In this case, a reduction in the base run length also results in a reduction of the colonization efficiency as cells fail to navigate the nutrient gradient in close proximity to the particle. In summary, the ability for bacterial cells to swim is fundamental to efficiently colonize particles at any settling velocity and chemotaxis plays an important role at intermediate settling velocities (10 m d^-1^) where the flow around the particle creates steep substrate gradients (in comparison to low settling velocities) but there remains sufficient time for the chemotactic behavior to navigate the particle boundary layer (in comparison to higher settling velocities where bacterial cells are rapidly swept around the particle).

### Particle microstructure facilitates colonization by non-motile cells

We extend our modeling approach to simulate scenarios that are infeasible experimentally and more representative of heterogeneous marine particles and aggregates in the ocean. Geometries that support intra-particle advective fluxes may readily emerge through the process of aggregation of smaller particles due to differential settling velocities or following polymer lysis by actively degrading bacterial colonies that colonized the particle previously. We craft a cohort of 500 stochastically generated particle geometries ranging from small (100 µm diameter) to large (3 mm diameter) particles with target porosities between 0 (densely packed particles) and 0.7 (flocculent particles/aggregates). Each particle is composed of stochastically arranged sub-particles that span between 10 µm and 100 µm diameter and permit interstitial advective flows (Fig. 4). We calculate the macroscopic settling velocity for each aggregate as a function of overall particle size and porosity (Methods). By using an individual based approach, we can relate the initial location and exact trajectory of each bacterium to the ability of the cell to find the particle. We used a larger domain for the porous particle simulations to ensure that there are no boundary effects on the flow around the particles. We validate that this does not significantly change our results by comparing the large domain and experimental twin geometry for a 3 mm solid particle at the same simulated velocities used in the experimental results (SI Fig. 3, SI Table 2). The model predicts a lower colonization efficiency for the large domain at slow settling velocities (1 m day^-1^) where the bacterial swimming velocity is greater than the flow velocity (and cells swim away from the particle) whereas no difference is predicted at higher settling velocities. Since all porous particles are simulated with a settling velocity greater than 20 m day^-1^, the difference in simulated domain geometry does not bias the colonization results (SI Table 2).

**Figure 4:**
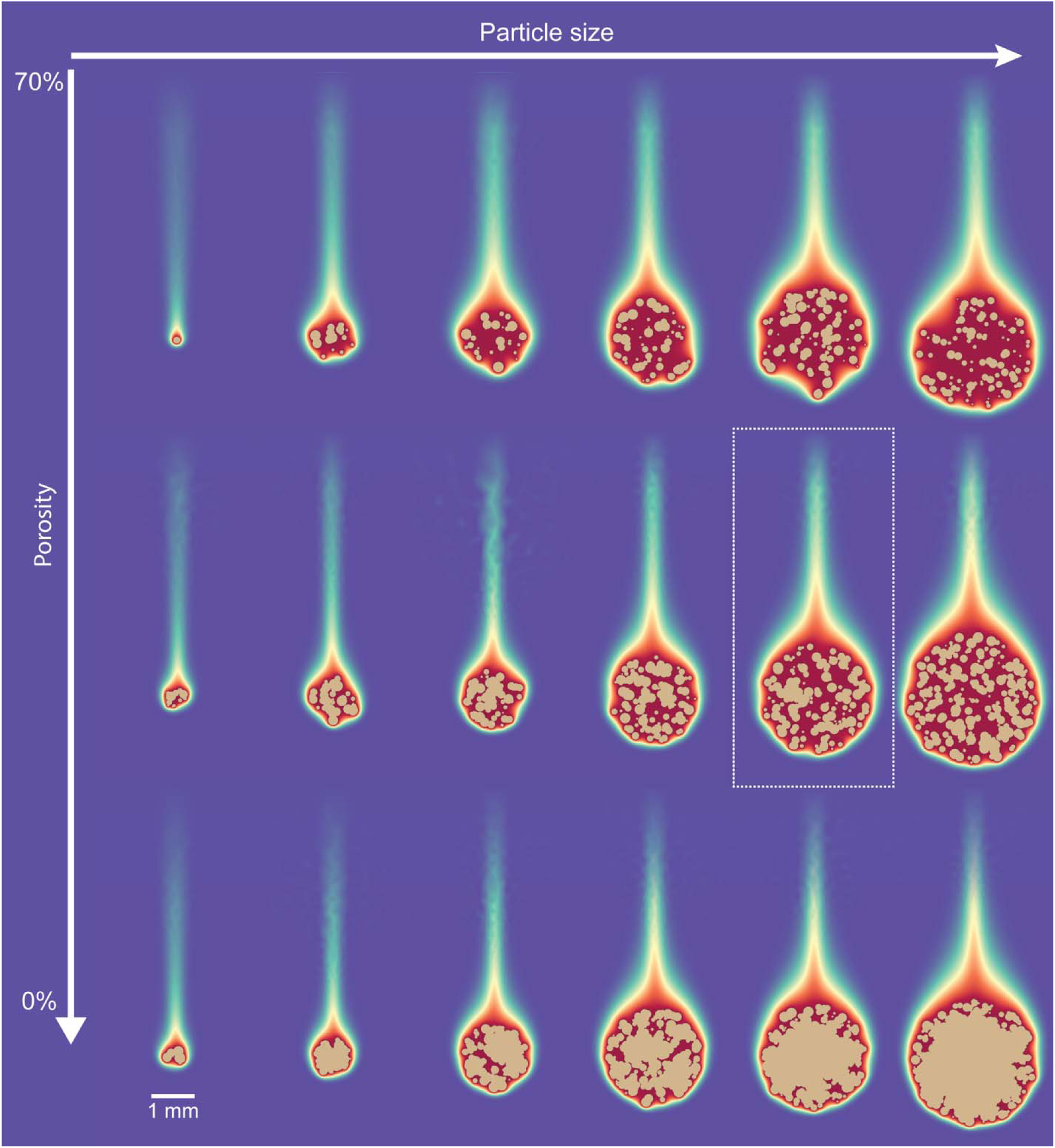
Example stochastic particle geometries and concentration fields. The density and settling velocity of the stochastic particles is determined by the number of aggregated sub-particles and overall particle size. These parameters further govern the resulting particle plume which is visualized as the predicted concentration field around the particle. The localized flow field, nutrient concentration and colonization by bacterial cells of the emphasized particle is highlighted in Fig. 5a.

Interestingly, the capture radius of the stochastically generated particles for both the chemotactic and motile cells is a function of the particle settling velocity and is notably governed by bacterial random walk motility rather than chemotactic behavior as might be expected *a priori* (Fig. 5b). The capture radius is defined as the radial distance from the stagnation line (the streamline colliding with the stagnation point from which the other stream lines diverge) that contains 95% of the cells that colonized the particle. This is expected since there are no nutrient gradients upstream of the particle for the chemotactic cells to navigate and further supported by the fact that we can predict the capture radius for both the chemotactic and motile cells via the characteristic diffusion distance for motile cells. For non-motile cells on the other hand, the observed capture radius is always larger than predicted by diffusive theory, indicating that they benefit disproportionally from the particle porosity. A higher particle porosity increases the colonization efficiency for all strains (Fig. 5c) and further benefits the chemotactic strain when compared to the motile strain (Fig. 5d). Higher porosity results in less flow being diverted around the particle and, consequentially, a diminished boundary layer and velocity gradient. The chemotactic cells benefit disproportionally from this phenomenon as the lower flow velocity in close proximity of the particle permits ample time to chemotactically navigate the boundary layer. The relation between the colonization efficiency and additional particle properties is shown in SI Fig. 4.

**Figure 5:**
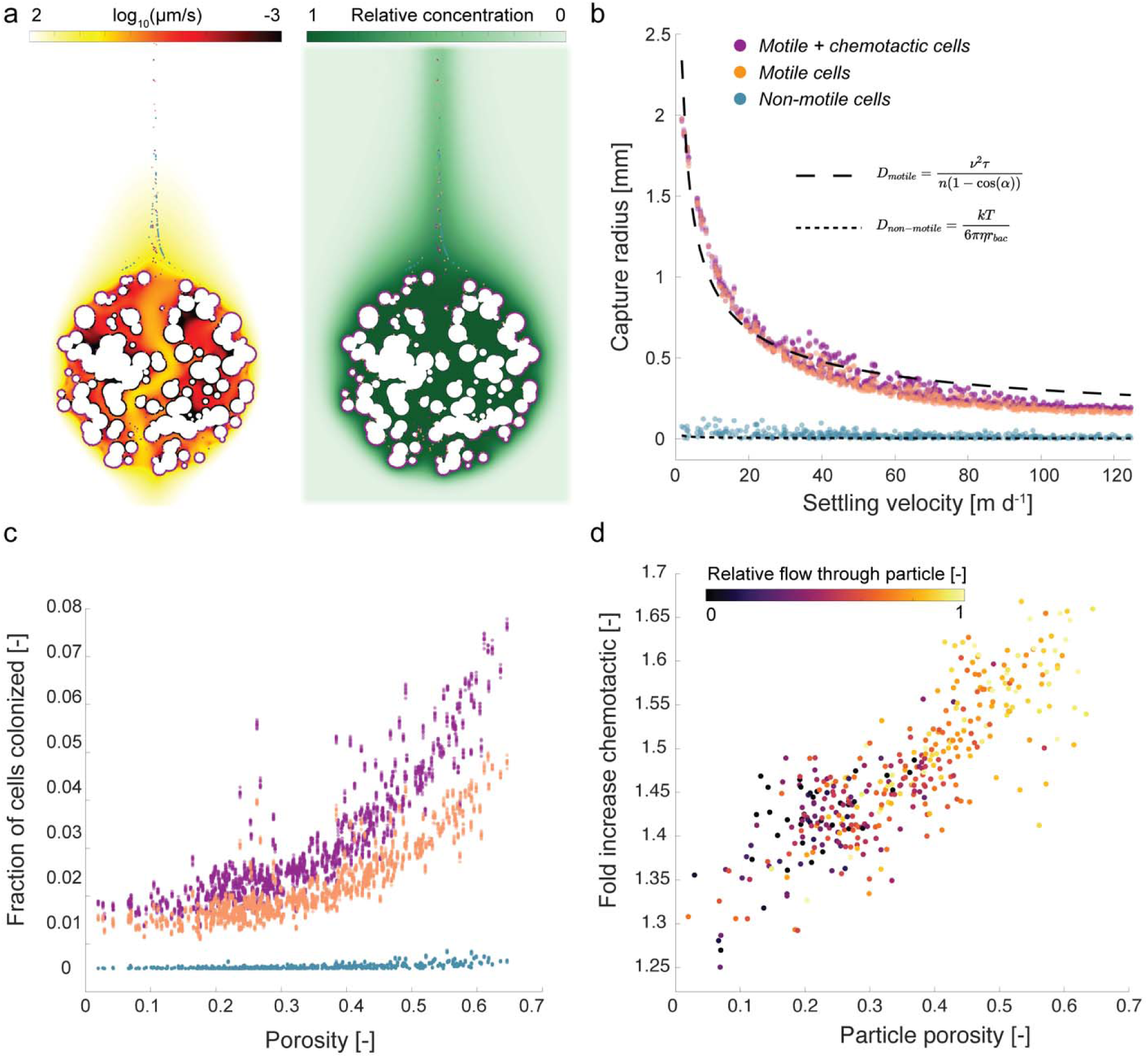
Quantifying bacterial colonization of porous particles. a) We use a cohort of 500 stochastically generated particles with varying sizes and porosities to simulate more realistic particle geometries. Porous particles have a similar macroscopic distribution of nutrient around them but permit interstitial advective flows that change the localized flow field around the particles. b) Capture radius of the particles as a function of their settling velocity. The capture radius is the radial distance from the stagnation line that contains 95% of the cells that colonized the particle. Results agree well with the characteristic diffusion distance calculated as 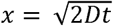 where t is the advective time from the inoculation point to the front of the particle and D the diffusivity of motile or non-motile cells. c) Fraction of the inoculated cells that colonize the particle as a function of the particle porosity. d) Increase in colonization by chemotactic cells compared to motile cells as a function of porosity and total flow through the particle.

To distinguish between the effects of particle porosity, roughness, and a smooth surface, we created two additional twin geometries for each of the 500 stochastic porous particles that use the same sub-particle structure with a filled core (particle with rough surface) or filled core and smoothed surface (Fig. 6b inset). The chemotactic and motile strains generally show a 10-fold increase in colonization efficiency when colonizing a porous compared to a rough particle (Fig. 6a). In comparison, the non-motile strain shows a far wider distribution dependent on the specific particle shape (log-ratio of 0) whereas others show a fold-increase of more than 1000. A similar trend is observable when comparing the rough to the smooth geometry (Fig. 6b). Both the chemotactic and motile strains show an overall negligible increase in colonizing a smooth virus rough “solid” particle whereas the non-motile strain shows a far more pronounced benefit with some particle geometries resulting in a 100-fold increase in colonization efficiency. However, despite having the greatest relative benefit from particles being porous or rough, the overall number of non-motile cells on the particle remains negligible compared to chemotactic or motile cells (Fig. 5c). The significantly higher colonization rate of the chemotactic and motile strains also has consequences for their relative abundance in the particle plume (Fig. 6c). The composition of a homogenized community (i.e., exact same relative abundances of the three strains) shifts downstream of the particle due to the differential colonization efficiency, resulting in an enrichment of non-motile cells in the particle wake. This enrichment is particularly amplified at slow settling velocities, representative of the vast majority of marine particles^37^. Despite not making it onto the particle, these non-motile cells play a crucial role in particle-associated nutrient cycling as their cumulative time in high concentrations of labile carbon (i.e., in close proximity to the particle or residing within the particle plume) is prolonged due to their residence in slow-velocity areas in the particle boundary and following the shift in relative abundance in the particle plume (Fig. 6d).

**Figure 6:**
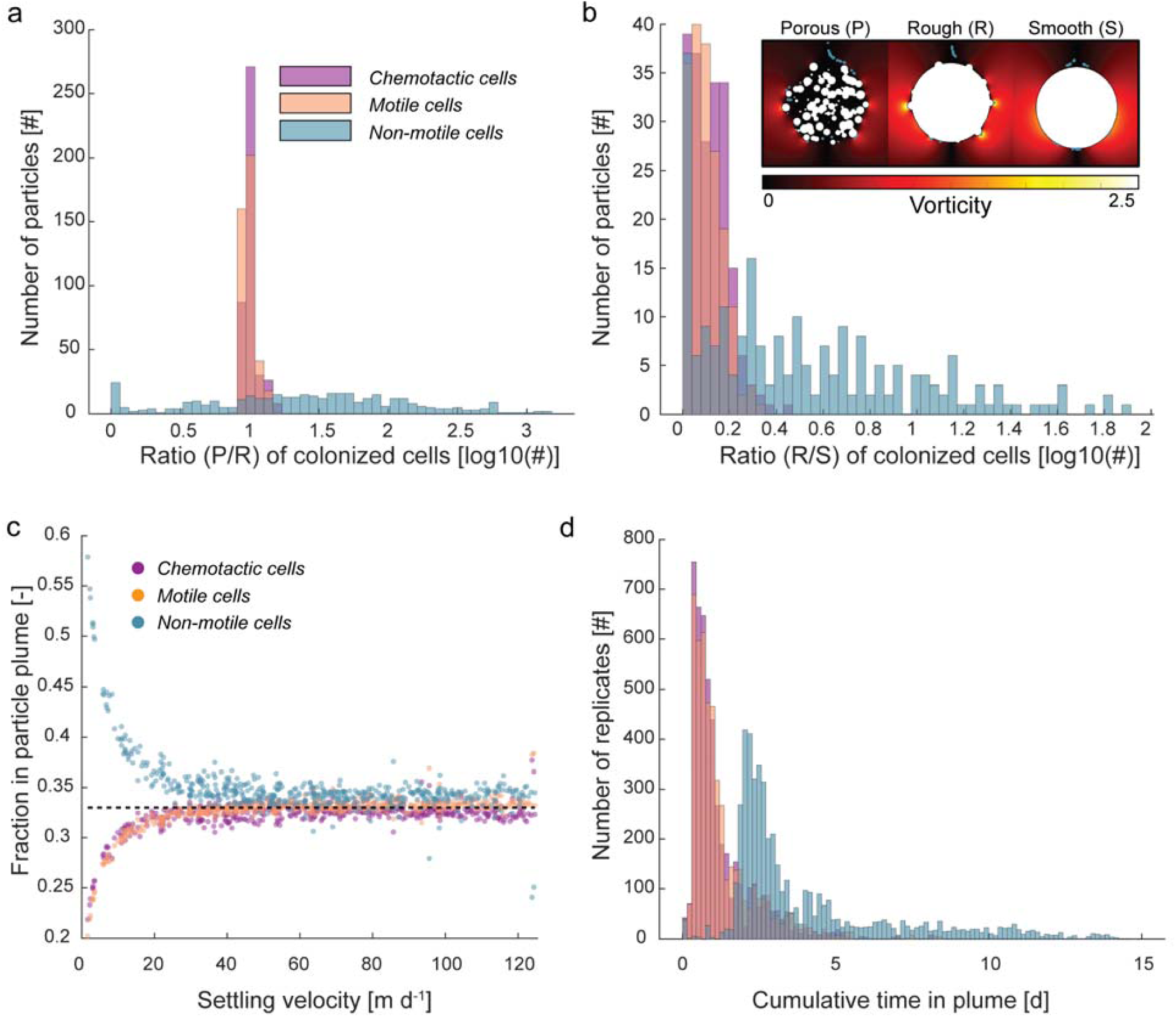
Influence of the particle micro-structure on the colonization efficiency and resulting community dynamics in the particle plume. a) Ratio of cells that colonized the porous particle (P) versus the rough particle (R) (see inset in panel b for an example comparison of the three geometries). b) Ratio of cells that colonized the rough particle (R) versus the smooth particle (S) for all species. c) Composition of the community aft of the particle as a function of settling velocity for a community with equal proportion of the strains upstream of the particle (dashed line). d) Distribution of the cumulative time (sum of the time spent within the plume across all cells for a replicate) across all particles.

## Discussion

### The benefit of chemotaxis and motility for bacterial colonization of marine snow

In this study, we visualized and quantified the colonization of marine particles by bacteria with different motility traits and highlighted the important influence of particle micro-structures on the colonization efficiency. Our experimental system enabled direct quantification of the colonization of a synthetic leaking particle by bacterial strains with different motility traits using fluorescence microscopy. A key improvement of this experimental system is the ability to mimic a leaking particle over a time period of multiple hours with controllable settling velocities. This enables us to observe how chemotaxis and motility facilitate the colonization of marine particles under realistic flow and chemoattractant conditions that was not possible with previous experimental approaches. Furthermore, by using mutants of *P. aeruginosa* (in contrast to different motile and non-motile marine species) and tagging the strains with the same fluorescent protein, we can directly compare and isolate the influence of the different motility traits on the colonization efficiency across the settling velocities. Our experiments and simulations clearly show that motility is required to colonize particles efficiently, in line with previous experimental results that quantify the colonization of curved surfaces by bacteria^17^. Our results show that chemotaxis indeed further benefits the bacterial cells to rapidly navigate the particle boundary layer, especially at higher settling velocities (Fig. 2a,b). The early colonization of marine snow is especially important for rapidly sinking particles. The low colonization efficiency and high growth rate of the copiotrophic species that typically find a particle favor a scenario where a few lucky cells establish themselves and rapidly proliferate, resulting in a “local endemism” that facilitates the emergence of a low particle-associated diversity and dominance of only a few taxa^38^.

Choosing a single, tractable species for the experiments does not fully elucidate the breadth of diverse motility and chemotaxis mechanisms found in the oceans. Although *P. aeruginosa* is not a typical marine bacterium (albeit one found in the ocean^39,40^), it is a model organism to study the chemotactic and motile behavior of singly flagellated cells that are representative of the majority of the marine species^41^. While *P. aeruginosa* (swimming velocity of 45 to 60 µm s^-1 17,42^) is considered a rapid swimmer, motile marine bacteria can be even faster, frequently exceeding speeds over 100 µm s^-1^ and potentially reaching up to 445 µm s^-1^ when tracking nutrient patches^43^. Indeed, bacterial swimming speed has previously been designated a key parameter for efficient chemotaxis, with an optimum as a function of the nutrient patch size^44^. Bacterial motility in our simulations is modelled as a classic run-and-tumble algorithm with a uniform turn angle distribution to minimize bias to the computational results due to parameter choice. However, *P. aeruginosa* motility differs and follows a “run-reverse” or “run-reverse-pause” pattern with a different turn angle distribution^45^. We strategically tested the influence of different turn angle distributions on the resulting colonization efficiency of the experimental twin geometry and found that the trend of reduced colonization efficiency as a function of the settling velocity and higher colonization efficiency by the chemotactic cells remains unchanged (SI Fig. 5). To further explore the importance of different aspects in a bacterial run and tumble movement (such as the average run length, swimming velocity or chemotactic sensitivity) to colonize the marine particles, we strategically explored the parameter space of the motility algorithm in the individual based simulations. Interestingly, the bacterial swimming speed has the greatest influence on the colonization efficiency in contrast to a change in the average run length that only marginally affects the colonization efficiency (Fig. 2b,3). This should result in an advantage for cells that evolve faster swimming speeds when relying on particle associated nutrients. For the parameter space exploration, we varied both the swimming velocity and chemotactic sensitivity over two orders of magnitude whereas we only changed the average run length over one order of magnitude to remain within a physiologically meaningful realm (e.g., average run lengths of 100 s are unrealistic). Contrary to our initial expectation, varying the chemotactic sensitivity across two orders of magnitude had a lesser influence on the colonization efficiency when compared to the swimming speed (Fig. 2b). We attribute this to the fact that bacterial cells already need to be in close proximity of the particle to sense the chemoattractant and the additional benefit of chemotaxis primarily comes into play when navigating the velocity gradient to the particle surface. Evidently, the chemotactic response of marine bacteria has to be very efficient in order to benefit from the brief time window where bacterial cells are within a nutrient gradient of a particle. Indeed, when compared to chemotaxis in *E. coli* (representing the most studied chemotactic system), marine bacteria show far more rapid chemotactic responses^27,46^ and other adaptions such as a velocity dependent motile behavior^46–48^ that suggest an evolutionary adaptation to profit from these ephemeral opportunities.

Finally, even when getting to the surface of the particle, bacterial strains show different strategies for surface attachment beyond the dynamics explored here. For example, strains as closely related as *P. aeruginosa* PAO1 and PA14 show stark differences in their strategy to surface attachment and detachment^49^. Even within the same bacterial population, the actual surface attachment strength cannot be deduced from population average observations as there is significant variability at the single cell level^50^. These observations preclude directly translating our specific quantitative observations to the dynamics of particle attachment and detachments in the real ocean beyond the initial encounter. For this reason, we primarily focus on the initial encounter and attachment of the bacterial cells in the model (perfect sticking condition) and observe the bacterial cells after a fixed amount of liquid has flown past the particle in the experiments to assess the influence of the initial colonization of the particles rather than the subsequent dynamics that emerge due to differential attachment and detachment. As such, our results describe the maximum potential of colonization. Nevertheless, the initial encounter and attachment of bacterial cells to the particle is of fundamental importance since it is the first and necessary step of planktonic cells to colonize marine sinking particles, contribute to the remineralization of marine snow, and thus influence the biological carbon pump efficiency as a whole.

### Particle heterogeneity and microstructure benefits non-motile cells

The geometry and structure of idealized disk-shaped synthetic particles is in stark contrast to natural particles that can be highly heterogeneous at the micro-scale^13,51,52^. In contrast to perfect spheres, the irregular shape creates steep velocity gradients and heterogeneous flow fields close to the particle surface^53^ that affect the bacterial trajectory during colonization. In addition, the flocculent and porous appearance of natural marine particles advocates the presence of interstitial advective flows that permeate the particles. Previously, a discrepancy between the predicted diffusive oxygen flux and experimentally observed oxygen consumption rates suggested the presence of advective oxygen flux into the particles and interstitial advective flow on the order of 40 µm s^-1 54^. Permeation of marine snow profoundly changes the way these settling particles interact and scavenge smaller particles in their path with scavenging efficiencies that are 100 times higher for porous particles compared to their smooth counter parts, even when deflecting most of the flow around the particles^55^. Interestingly, the majority of this scavenging is attributed to a thin boundary layer just interior to the particle surface. This is critical since the process of scavenging is therefore not dependent on total perfusion of the particle^55^ and can still occur if the center of the particle restricts advective flows due to, e.g., clogging of the interstitial pore space by transparent exopolymers (TEP)^56^. These theoretical considerations are in good agreement with our experimental observations where a higher particle porosity facilitates the colonization by all three strains (Fig. 4c). In summary, permeability of marine snow (even if restricted to the surface) can play a principal role in shaping the colonization by chemotactic and motile cells (by reducing the total deflection of flow around the particles and reducing the velocity gradients), but primarily facilitates some colonization by non-motile cells by allowing stream lines to intercept with the particle surface and thus passively increases the encounter between non-motile cells and marine snow.

Since natural marine snow is often associated with the presence of transparent exopolymers (that permit diffusive fluxes but diminish advective flow), we further investigate how only surface roughness changes colonization of bacteria with different motility traits (Fig. 5). Except for a few particle geometries where the stochastic geometry favors rapid flushing of the bacterial cells through the particles, porosity primarily benefits the non-motile cells when calculating the fold increase in colonization between the rough and porous simulations (Fig. 5a). Similarly, the non-motile strain showed the highest fold increase when comparing the colonization efficiency between the rough and smooth particles (Fig. 5b). The increase in colonization efficiency for the non-motile strain is mainly driven by the small-scale alterations of the flow field around the particle that enable non-motile cells that are in close proximity of the surface to attach to the particle. Recently, experimental observations using particle image velocimetry showed similar small-scale alterations of the flow field around particles of heterogeneous shapes when compared to spherical particles^53^. Combined with our results, these observations suggest that even in absence of any particle porosity that supports interstitial advective flows, the microscale heterogeneity of the particles can promote the colonization of non-motile cells.

Nevertheless, the colonization efficiency of non-motile cells is orders of magnitude lower when compared to both the chemotactic or motile cells, especially at lower settling velocities (Fig. 5c, SI Fig. 4). This may have profound consequences for the planktonic bacterial community composition in the bulk ocean (i.e., non-particle associated community). For the North Pacific, it is estimated that the average time between particle encounters for motile bacteria is on the order of tens of hours, but 10% of the bacteria find a new particle within a few hours and the top 1% encounters a settling marine particle within minutes^57^. The effective diffusivity of non-motile cells is orders of magnitude lower when compared to motile strains^58^ and the resulting encounter rate between particles and non-motile cells becomes vanishingly small. However, non-motile cells may still benefit from the rapid encounter of motile and chemotactic cells with the marine snow. In absence of additional cells from outside of the ballistic and/or diffusive kernel, we observe an enrichment of the non-motile cells in the plume of the particle (Fig. 6c), especially at slow settling velocities where the colonization efficiency of the motile and chemotactic cells is high. In this case, the particle effectively acts like a sponge that selectively scavenges chemotactic and motile cells from the local environment and exports them to the ocean interior. Additionally, despite having to share this ephemeral nutrient hotspot with chemotactic cells that migrate towards the plume^27^, non-motile cells can profit from the rapidly diffusing nutrient plume that benefits cells in a far larger area compared to the predicted encounter kernel. Since this radial diffusion promptly dilutes the nutrients within the plume, the non-motile cells that are frequently associated with an oligotrophic lifestyle are better equipped to benefit from these ephemeral but common sources.

The ability of bacterial cells to exploit the nutrient rich microenvironments presented by marine snow is a key process in global biogeochemistry. The encounter and attachment of bacteria to marine particles presents the first step of particle remineralization, and a mechanistic understanding of these processes is thus crucial to advance our predictive capabilities. Our study demonstrates the although motility and chemotaxis are fundamental to efficiently colonize marine particles, non-motile cells benefit disproportionally from the porous microstructure of marine snow and associated interstitial advective flows. Non-motile cells can persist by benefitting from the wakes of ubiquitous and transient particles. The rapid scavenging of chemotactic and motile cells by settling particles and subsequent export to the ocean interior additionally shapes the planktonic community by enriching non-motile cells in the relatively nutrient rich particle plume. Our results thus provide a mechanism to explain the apparent paradox that the ocean interior is a dilute nutrient-poor medium where cells rely on motility to find nutrient patches yet one numerically dominated by non-motile cells. Indeed, this mechanism of scrubbing out chemotactic and motile organisms may help to explain, for instance, how SAR11, the most abundant organism on Earth and a non-motile cell^1^ has achieved its ubiquitous dominance in the ocean. Disentangling the interplay between the preferential colonization by motile cells guided by chemotaxis and the benefit for non-motile oligotrophic bacteria in the planktonic community across the global oceans warrants continued study.

---

**SI Figure 1:**
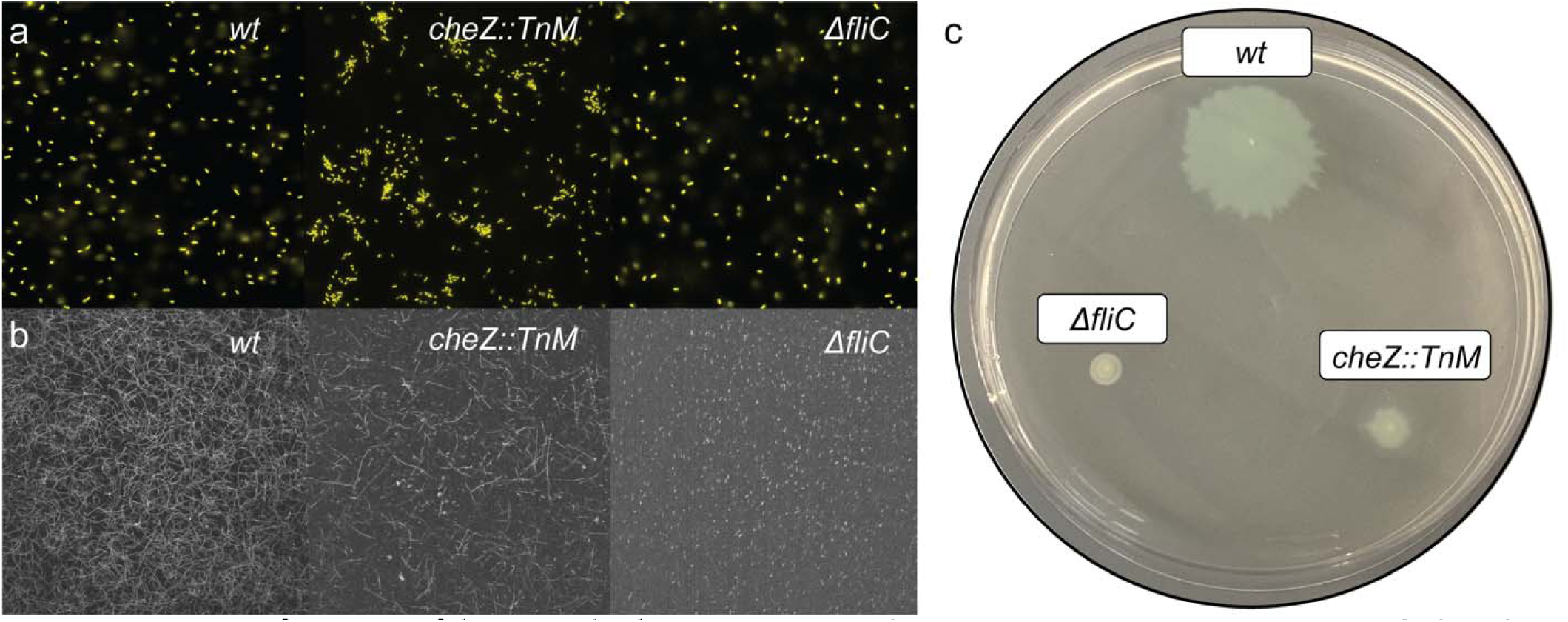
Verification of bacterial phenotypes. a) Fluorescence microscopy image of the three bacterial strains used in the study to verify the constitutive expression of the fluorescent protein. b) Bacterial swimming trajectories visualized using high-speed imaging. In contrast to the high percentage of motile cells for the WT and cheZ::TnM strains, the ΔfliC does not show any motile behavior at the single cell level. c) The motility and chemotaxis of the three strains was further verified with a soft agar approach^59^ that confirms the chemotactic deficiency but motile behavior of the cheZ::TnM strain and non-motile behavior of the ΔfliC mutant.

**SI Figure 2:**
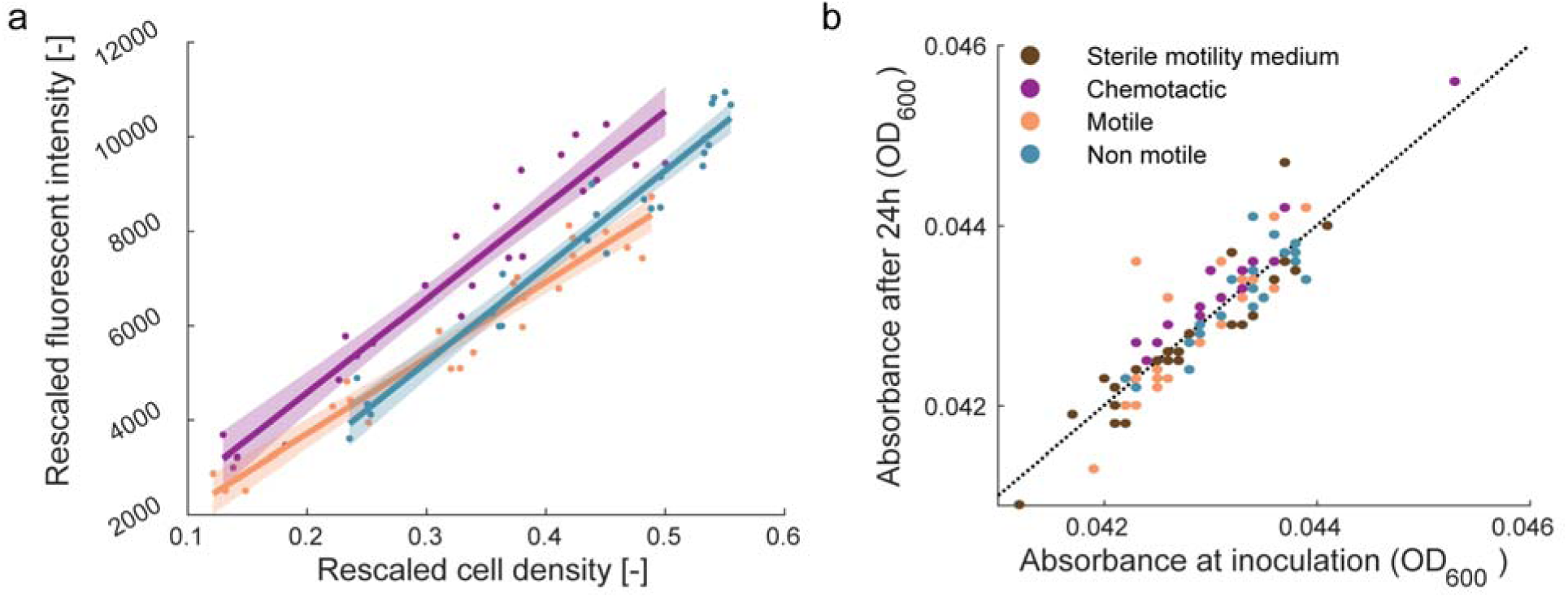
Density dependent bacterial fluorescence and growth in motility medium. a) Comparison of rescaled fluorescent intensity (measured intensity minus control intensity of sterile LB media) compared to the rescaled cell density (measurement of optical density at 600 nm minus the optical density at 600 nm of sterile LB media). b) Comparison of optical density of the different bacterial strains in motility medium containing each 10 µM L-serine and L-aspartate directly after inoculation at a density of 10^5^ cells per ml and after 24 h of incubation at 37 °C.

**SI Figure 3:**
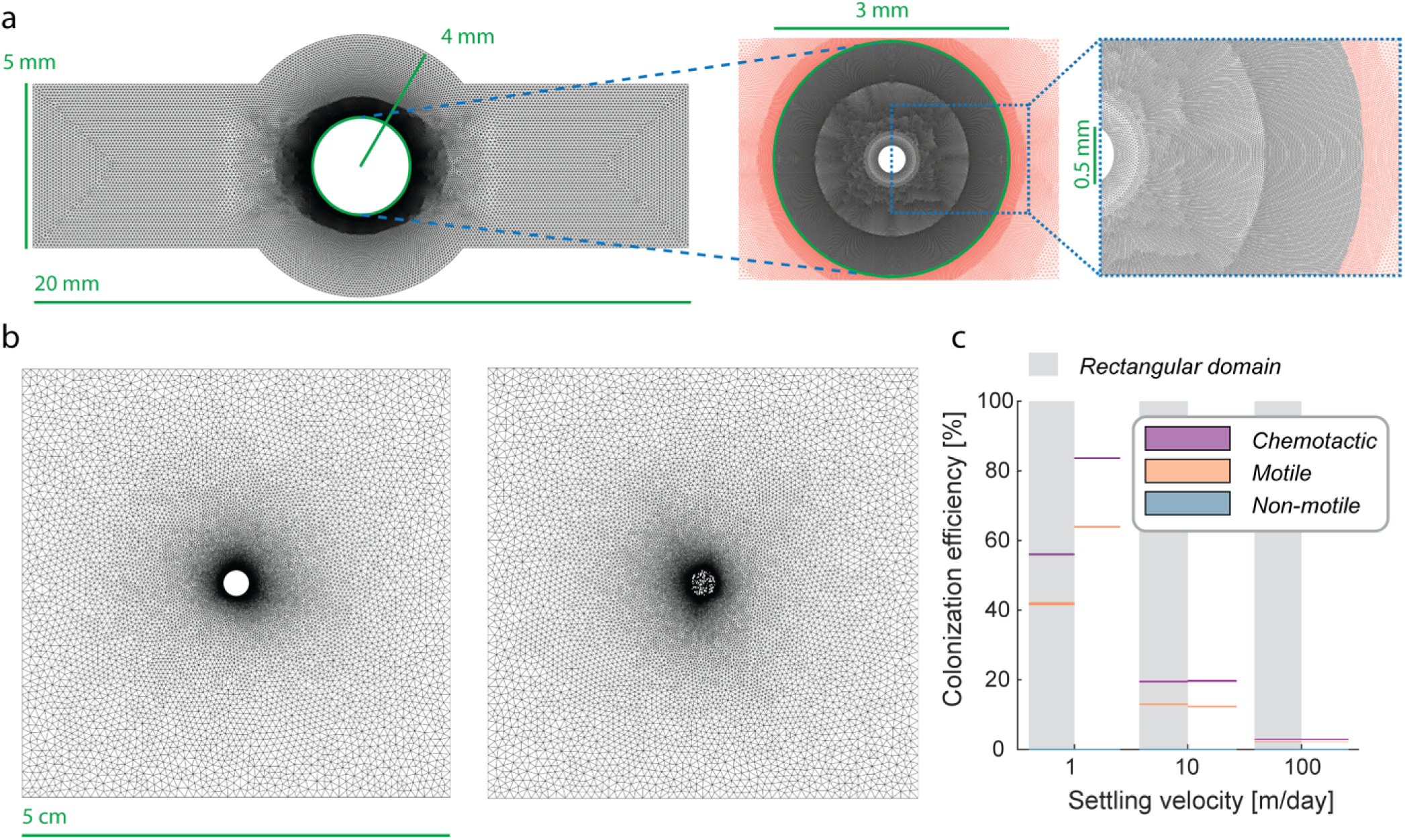
Visualization of in silico geometries of the different scenarios. a) The experimental twin scenario is represented by a 5 mm wide and 20 mm long rectangular channel including a 4 mm diameter expansion in the center that contains the 3 mm diameter particle. The grid mesh is 200 µm that increases in resolution to 10 µm resolution towards the particle boundary. To account for realistic radial diffusion, the whole particle is simulated at a 10 µm grid resolution with a constant carbon source of diameter 0.5 mm at the center (congruent to the experimental system). b) For the stochastic simulations of porous, rough and smooth particles, we increased the domain size to 5 cm to avoid any bias from potential boundary effects. We used a grid resolution of 1 mm that increases to 20 µm towards the boundary of the particles. c) The larger rectangular domain primarily shows lower colonization efficiency at low settling velocities when compared to the experimental twin system using the same flow velocities and inoculation conditions. This is due to bacterial cells diffusing away from the particle since they are not constrained by the flow and/or reflected from the boundary of the domain.

**SI Figure 4:**
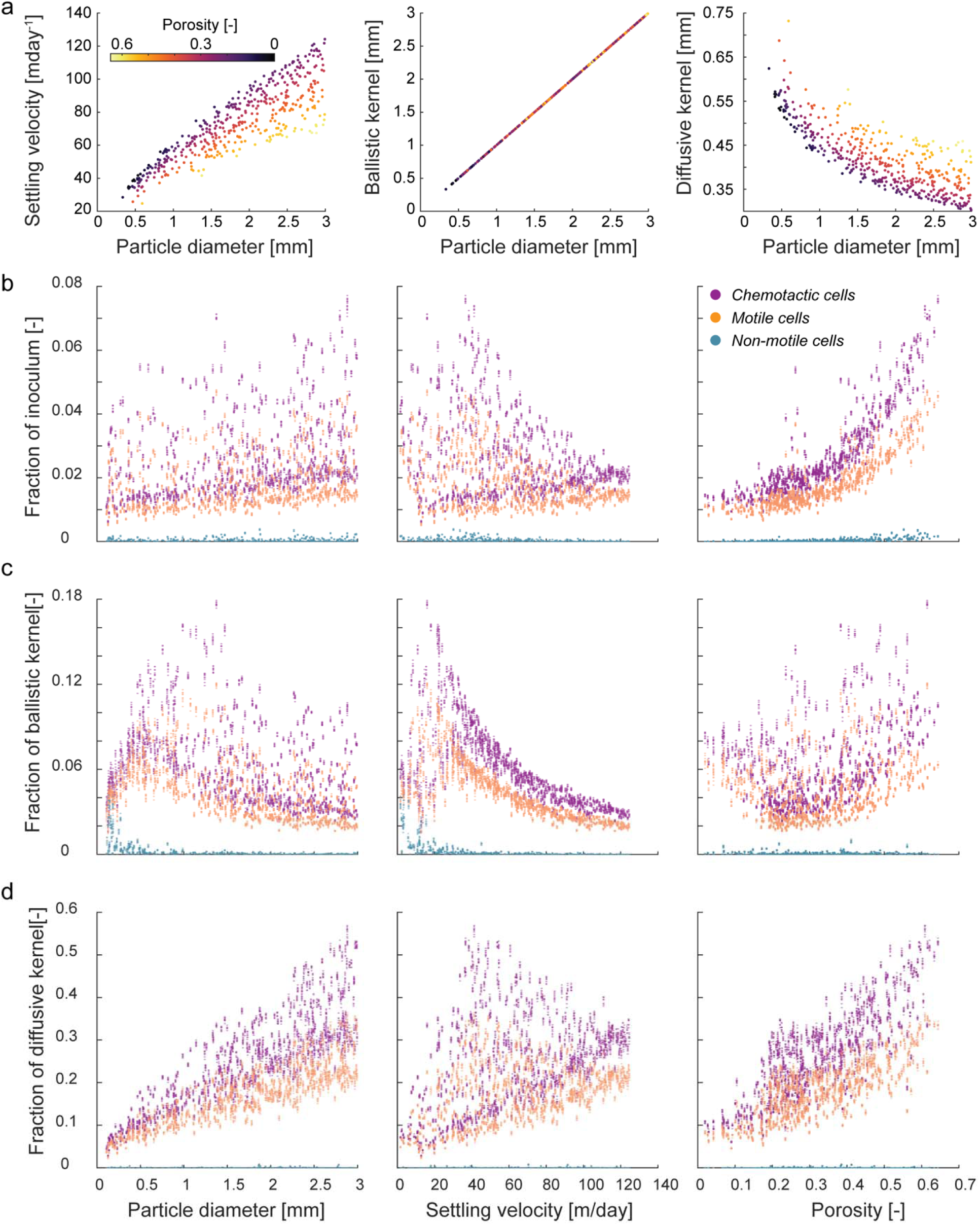
Comparison of different predictors to the colonization efficiency as a function of different encounter kernels. a) Relation between the different predictors (particle size, settling velocity and porosity), the diffusive kernel as a function of particle diameter and porosity and visual explanation of the different kernels. b) Relation between the colonization efficiency when taking into account the total inoculum and the different predictors. c) Relation between the colonization efficiency when taking into account the cells within the ballistic kernel and the different predictors. d) Relation between the colonization efficiency when taking into account the cells within the diffusive kernel and the different predictors.

**SI Figure 5:**
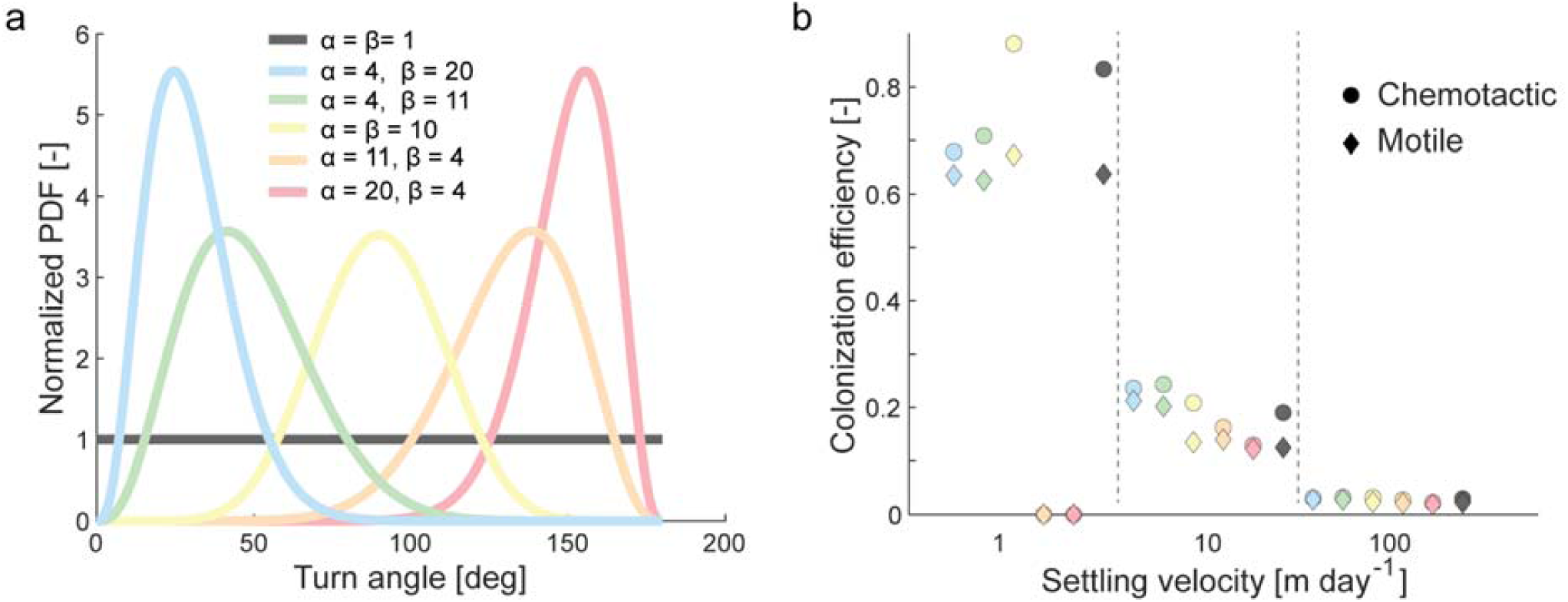
Exploration of different turn angle distributions on the colonization efficiency of chemotactic and motile cells. a) Simulations with varying turn angle distributions using combinations of the shape parameters *α* and *β* of a Beta distribution to simulate distributions ranging from uniform via Gaussian to left and right skewed distributions (see SI Appendix 1). b) Comparison of the colonization efficiency (N=10^5^ cells) for the different settling velocities and turn angle distributions. Chemotaxis has the highest influence at turn angle distributions with mean 90° (uniform and Gaussian). Left skewed distributions (orange and red) show no colonization at 1 m day^-1^ since the cell swimming velocity exceeds the settling velocity and frequent cell reversal results in an overall negligible particle encounter.

**SI Table 1:** Pairwise statistical comparison between and within the different strains for the experimental results. For all pairwise comparisons, we used two-sided t-tests. Comparisons were made between the different strains at the same settling velocities and within the same strain for different settling velocities. Statistically significant differences are shows in green whereas non-significant results are shown in red. Blue fields indicate comparisons that were not performed (between different strains at different velocities or with itself).

| Strain | m d<sup>-1</sup> | Chemotactic |  |  | Motile |  |  | Non-motile |  |  |
| --- | --- | --- | --- | --- | --- | --- | --- | --- | --- | --- |
|  |  | 1 | 10 | 100 | 1 | 10 | 100 | 1 | 10 | 100 |
| Chemotactic | 1 |  | 2.09E-04 | 7.05E-09 | 0.0213 |  |  | 4.80E-04 |  |  |
|  | 10 |  |  | 0.3084 |  | 0.19 |  |  | 0.1875 |  |
|  | 100 |  |  |  |  |  | 0.0066 |  |  | 0.0011 |
| Motile | 1 |  |  |  |  | 0.0103 | 0.0051 | 0.0364 |  |  |
|  | 10 |  |  |  |  |  | 0.2335 |  | 0.7781 |  |
|  | 100 |  |  |  |  |  |  |  |  | 0.9606 |
| Non-motile | 1 |  |  |  |  |  |  |  | 0.703 | 0.1102 |
|  | 10 |  |  |  |  |  |  |  |  | 0.5418 |
|  | 100 |  |  |  |  |  |  |  |  |  |

**SI Table 2:**
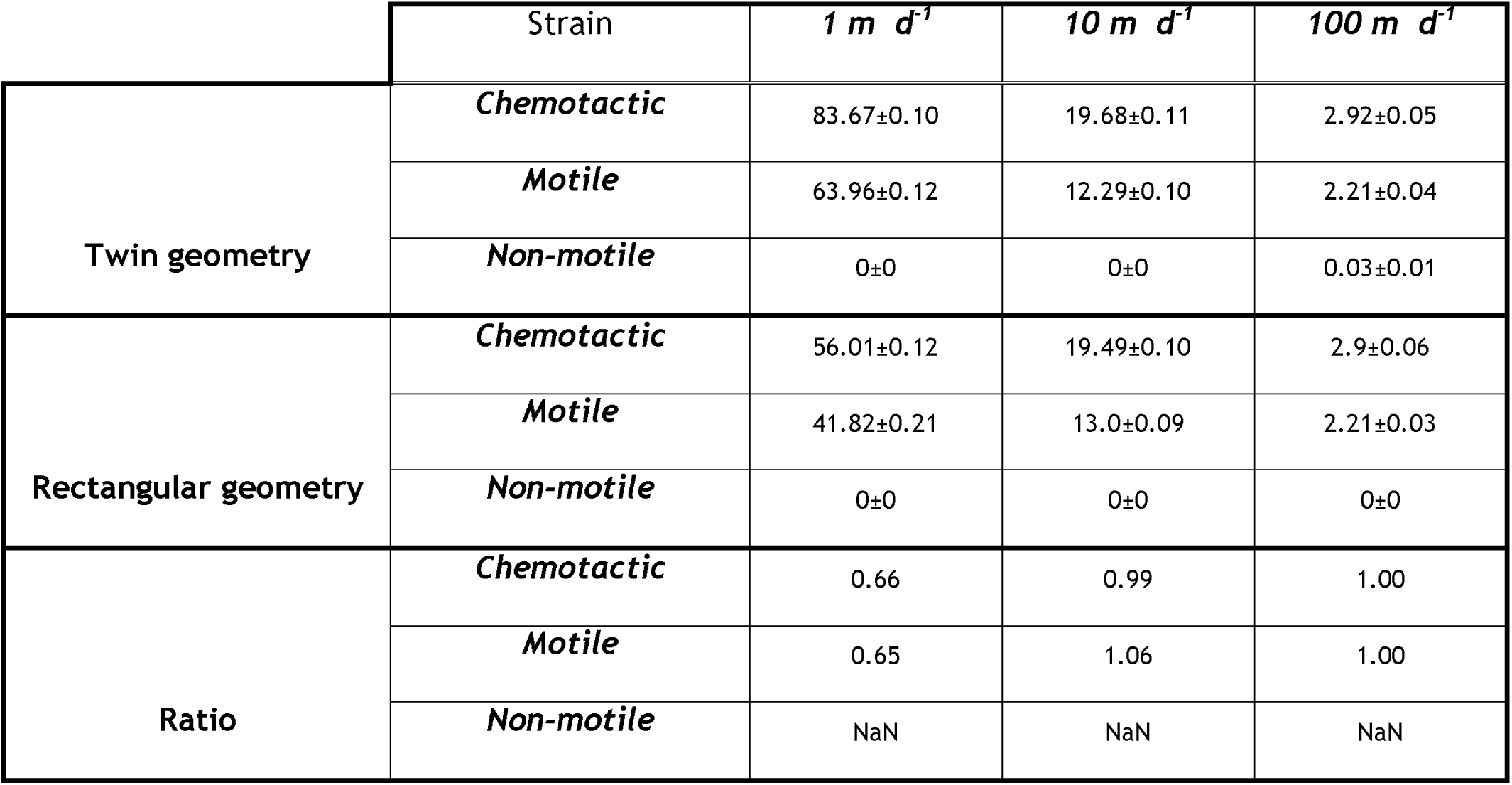
Comparison of in silico results for the experimental twin geometry and a large rectangular domain. Colonization efficiency (in %) for all strains and settling velocities in the simulations for both the experimental twin geometry and the rectangular geometry including the ratio of the colonization efficiency in the rectangular geometry compared to the experimental twin geometry.

**SI Table 3:** Mean, median and mode of the different turn angle distributions. Mean, median and mean turn angles are shows in degrees.

| $\alpha$ | $\beta$ | *Mean* | *Median* | *Mode* | *Type* |
| --- | --- | --- | --- | --- | --- |
| 1 | 1 | 90 | 90 | 90 | Uniform |
| 10 | 10 | 90 | 90 | 90 | Gaussian |
| 20 | 4 | 150 | 152 | 155 | Left skewed |
| 11 | 4 | 132 | 134 | 138 | Left skewed |
| 4 | 11 | 48 | 46 | 42 | Right skewed |
| 4 | 20 | 30 | 28 | 25 | Right skewed |

## SI Appendix 1

We explored the influence of different turn angle distributions on the resulting colonization using the numerical model where a new direction (angular turn) after tumbling is chosen at random from a Beta distribution. The Beta distribution is a combination of a power function of the variable x and its reflection (1-x) and is characterized by two shape parameters (*α* and *β* as shown in SI Equation 1 where Z is an arbitrary constant and B is the Beta function that normalizes the integral of the probability distribution function.

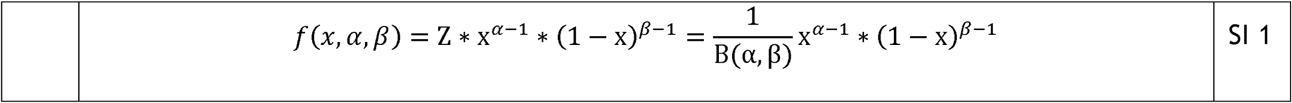

By strategically varying the two shape parameters, we can simulate different turn angle distributions from uniform, over non-skewed (Gaussian) to left or right skewed distributions (SI Figure 5). The mean, median and mode of the turn angles for the different combinations of the shape parameters are shown in SI Table 3. All simulations with varying turn angle distributions use the experimental twin geometries and all three simulated settling velocities.

